# Non-food crop *Rosa canina L* leaf and twig extracts as a source of nutrients and bioactive compounds

**DOI:** 10.1101/2020.04.06.027383

**Authors:** Małgorzata Kubczak, Ainur B. Khassenova, Bartosz Skalski, Sylwia Michlewska, Marzena Wielanek, Araylim N. Aralbayeva, Zhanar S. Nabiyeva, Maira K. Murzakhmetova, Maria Zamaraeva, Maria Skłodowska, Maria Bryszewska, Maksim Ionov

**Affiliations:** Department of General Biophysics, Faculty of Biology and Environmental Protection, University of Lodz, Lodz, Poland; Department of Biotechnology, Faculty of Food Production, Almaty Technological University, Almaty, Kazakhstan; Department of General Biochemisty, Faculty of Biology and Environmental Protection, University of Lodz, Lodz, Poland; Laboratory of Microscopic Imaging and Specialized Biological Techniques, Faculty of Biology and Environmental Protection, University of Lodz, Lodz, Poland; Department of Plant Physiology and Biochemistry, Faculty of Biology and Environmental Protection, University of Lodz, Lodz, Poland; Department of Biophysics and Biomedicine, Faculty of Biology and Biotechnology, al-Farabi Kazakh National University, Almaty, Kazakhstan; Department of Biophysics, Laboratory of Molecular Biophysics, Faculty of Biology and Chemistry, University of Bialystok, 15-245 Bialystok, Poland

**Author notes:** Author for correspondence: Dr. Maksim Ionov, Department of General Biophysics, Faculty of Biology and Environmental Protection, University of Lodz, 141/143 Pomorska Str., 90-236 Lodz, Poland.

**Keywords:** *Rosa canina L*, plant extract, bioactive compounds, cytotoxicity, antiradical activity

## Abstract

It is important to search for new sources of bioactive, natural compounds because customers pay more attention to food quality. Fruits and berries from horticultural plants are known to be good sources of agents beneficial for human well-being and could serve as natural preservatives in the food industry. However, more recent research indicates that other plant organs can also be rich in nutrients. Our study focused on characterizing an unexplored source: leaf and twig extracts from *Rosa canina*. The chemical composition of these extracts was analyzed and their *in vitro* activity measured. HPLC analysis of the content of phenolics, vitamins and amino acids revealed that the leaf and twig extracts are rich in bioactive compounds with potent antioxidant properties. The greatest differences between bioactive phenolic compounds in leaf and twig extracts related mainly to *p*-coumaric acid, myricetin, ellagic acid, cyanidin, procyanidin and quercetin, whereas salicylic acid levels were similar in both types of extract.

Interactions with human serum albumin were investigated and some conformational changes in protein structure were observed. Further analysis (lipid peroxidation, protein carbonylation, thiol group oxidation, DPPH inhibition and ROS inhibition) confirmed that both leaf and twig extracts exhibited antioxidant and antiradical scavenging activities. Cytotoxicity and hemotoxicity assays confirmed very low toxicity in the extracts over the range of concentrations tested. Our results indicate that both extracts could serve as non-toxic sources of bioactive compounds with antiradical properties.

## Introduction

The relationship between balanced diet and human health is well documented [1,2]. Moreover, knowledge of the beneficial effect of a diet enriched in fruits, vegetables, herbs and wild plants as rich sources of natural compounds with anti-oxidative, anti-inflammatory, anti-bacterial, anti-diabetic and anticancer properties is general accessible [3–7]. However, only a select group of customers pay particular attention to food quality. For most people, the everyday diet generally comprises highly transformed and manufactured food products that are rather poor in many vitamins, minerals and other compounds beneficial for health. On the other hand, today’s consumables markets offer products from new food categories called ‘functional food’. These products are enriched in compounds beneficial for health originating from plants, many of which are well known and have been used for thousands of years. However, the organs of these plants that have been available for years are now being used for technological processes, whereas other organs from them could be more interesting sources of beneficial compounds and more valuable for human consumption.

*Rosa canina* L. belongs to the *Rosaceae* family, which contains more than 100 species, and grows mostly in Europe, Asia, North America, Africa and the Middle-East [8]. *R. canina* pseudo-fruits (hips) are the best characterized organ of this plant. The hips were used worldwide as an antioxidant, anti-inflammatory, immunosuppressive, cardioprotective, gastroprotective or antimicrobial agent [9,10]. Nowadays, hip extracts are commonly used in and the cosmetic and food industries [11,12].

Many publications indicate that rose hips contain large amounts of vitamins A, B, C, D, and E, minerals, carotenoids, and phenolic compounds [10,13–17]. They also contain fruit acids, pectin, sugars, organic acids, amino acids and essential oils [18]. Vitamin C and phenolic compounds are well known for their antioxidant properties [7,8,14,19,20]. *R. canina* hips contain the highest level of the *L*-isomer of vitamin C among fruits and vegetables [21,22]. The ascorbic acid content of rose hips ranges from 300 to 4000 mg/100 g, the variation resulting from changes in sugar levels during ripening [16].

Vitamin E is another strong antioxidant. The human diet should include components rich in tocopherols. The lipid-soluble vitamin E is necessary for different antioxidant functions in human cells, especially in cell membranes and plasma lipoproteins. It helps to prevent the proliferation of oxidative chain reactions by scavenging many reactive oxygen species and it could be implicated in the prevention of atherosclerosis and cancer. Epidemiological investigations have revealed a positive correlation between tocopherol intake and a reduced risk for cardiovascular diseases [12,23].

Phenolic compounds confer the unique flavours and health-promoting properties of vegetables and fruits [24]. Coloured rose fruits are good sources of phenolic compounds including bioflavonoids, tannins, flavonoids, phenolic acids, anthocyanins and dihydrochalcones [25]. These compounds have a wide spectrum of biochemical activities such as antioxidant, antimutagenic, anticargionogenic effects and can alter gene expression [26]. Flavonoids and phenolic acids have diverse positive biological activities, making them the most important groups of secondary plant metabolites and natural bioactive compounds for humans [27].

HPLC analysis has shown *R. canina* hips extract to be especially rich in the polyphenols hyperoside, astragalin, rutin, (+)-catechin and (-)-epicatechin, gallic acid and poly-hydroxylated organic acids such as quinic acid [28,29]. The phenolic compouds (+)-catechin, (-)-epicatechin, rutin, vanillin, astragalin, phloridzin and gallic acid, identified in hips extract, have been reported as strong scavengers of the ROO^-^ radical [30]. Several other polyphenols potentially beneficial for humans such as ellagic acid, salicylic acid, vanillic acid, ferulic acid and caffeic acid have been identified in trace amounts in *R. canina* hip extracts [31,32].

Food products containing *R. canina* compounds are derived from the hips [33]. The leaves and stems from the roses are usually discarded as trash. However, increasing numbers of publications confirm that *R. canina* leaves could be a valuable source of flavonoids, especially flavone glycosides [34–36]. A few recent studies have indicated that stems from *R. canina* are also a good source of polyphenols [37,38]. Ouerghemmi et al. showed that stem extracts from different *Rosa* species could be used in the food, cosmetic and pharmaceutical industries as a source of phenolic compounds.

The aim of our study was to determine the biological properties of extracts of *Rosa canina* twigs and leaves. First, we focused on identifying compounds such as phenolics, amino acids and vitamins in the extracts. Secondly, we assessed the antioxidant and antiradical properties of these extracts. Also, the interaction of the extract components with human serum albumin was tested by circular dichroism and the hemotoxicity and cytotoxicity of the extracts were determined.

## 2. Materials and methods

### 2.1. Plant material

The full expander (development) leaves and twigs of *Rosa canina* L. were collected from the foothills of the Trans Ili Alatau Mountains (Almaty region, Kazakhstan) during the early morning in June 2018, in the middle of the vegetative season. The samples were obtained 1.5-2.0 meters from the ground, 5-8 cm from the branch meristem, from 10-15 year old plants. The plants were identified and voucher specimen No. 3389 (*R. canina* L.) was deposited at the herbarium of the Institute of Botany and Phytointroduction (Almaty, Kazakhstan).

### 2.2. Preparation of plant extracts

The *R. canina* leaves and twigs were thoroughly washed with distilled water and dried at room temperature, then crushed and extracted with 50% ethanol for 20 hours at 25 ± 2°C on a rotary shaker (110 rpm). After centrifugation (20 min, 20,000 rpm) the supernatants were dried under vacuum using a rotary evaporator at 50°C. The evaporated extracts were stored at 4°C. Before analysis they were dissolved in distilled water.

### 2.3. HPLC analysis

The HPLC system (Dionex, Sunnyvale, USA) was equipped with a photodiode-array detector. The compounds were separated on a RP column (aQ Hypersil GOLD, 250 × 4.6 mm, 5 µm) joined with a guard column (GOLD aQ Drop-In guards, 10 × 4 mm, 5 µm, Polygen, Gliwice, Poland) at 25°C using a mobile phase composed of water (A) and methanol (B), both with 0.1% formic acid. The linear gradient was started after 2 min of isocratic elution with 5% B, increasing slowly over 30 min to 55% B, followed by 5 min of isocratic elution, then an increase to 70% within 10 min; then, after an isocratic step with 70% B for 5 min, the gradient was returned to the initial 5% B within 2 min to re-equilibrate the column for the next 3 min. The flow rate was 1 cm^3^ min^-1^, and the absorbance was measured at 210, 235, 280, 325 and 375 nm.

Compounds in the *R. canina* extracts were identified by comparing the retention times and on-line UV absorption spectra of the analysed samples with the respective data obtained from reference standards. Quantification was based on a calibration curve for standards covering the range 5-200 µg cm^-3^; the linearity of the calibration curve was verified by the correlation coefficient (r^2^=0.9994).

### 2.4. Determination of phenolic compounds

The phenolic content was determined using Folin–Ciocalteu reagent according to the Singelton and Rossi (1965) method [39]. The absorbance of the reaction product was measured at 725 nm and the phenolic content was expressed as milligrams per gram of dried extract based on the calibration curve prepared for chlorogenic acid (Sigma-Aldrich). The results are given as means ± SD (n = 3).

The flavonoid content was determined by the aluminum chloride colorimetric method according to Chang et al. (2002) [40]. The absorbance of the reaction mixture was measured at 415 nm and the flavonoid content was expressed as milligrams per gram of dried extract based on a calibration curve prepared for quercetin (Sigma-Aldrich). The results are given as means ± SD (n = 3).

The catechin (flavan-3-ol) content was determined by the vanillin assay method described by Bakkalbasi et al. (2005) [41]. The absorbance of the reaction mixture was measured at 500 nm and the total flavan-3-ol content was calculated from a calibration curve prepared using (+)-catechin (Sigma-Aldrich) and expressed as milligrams per gram of dried extract. The results are given as means ± SD (n = 3).

### 2.5. Determination of water-soluble vitamins

A Kapel-105M Lumex (Russia) capillary electrophoresis kit was used to determine the vitamin composition of the extracts from leaves and twigs. The contents of B1 (thiamine), B2 (riboflavin), B3 (pantothenic acid) and B5 (nicotinic acid) were determined. The vitamins were detected at 200 nm and by using programmable wavelength switching. Conditions for separation: borate buffer pH = 8.9, temperature 30°C. The method was based on the extraction of free forms of vitamins and separation and quantification of the components by capillary electrophoresis.

### 2.6. Determination of tocopherol isomers

Isomers of vitamin E were identified by HPLC with UV detection. A high-performance Agilent 1200 chromatographic chromatograph (USA) with a four-channel thermostat pump, a spectrophotometric detector and a 250×4.6 mm Zorbax 300SB-C18 column was used. To determine the vitamin E content the following conditions were selected: flow rate of the mobile phase = 0.7 ml min^-1^; column temperature = 35°C. The eluant was 42: 50: 8 acetonitrile: isopropanol: water.

To prepare the samples they were extracted with an organic solvent, and then proteins that would interfere with the chromatography were precipitated. The samples were then evaporated to dryness and dissolved in 1 ml of the mobile phase.

### 2.7. Determination of amino acids

The Kapel-105M Lumex capillary electrophoresis (Russia) was also used to assess the amino acid composition of the extracts. The electric field separated the charged components of the extracts in a quartz capillary. A microvolume of the solution to be analyzed (∼ 2 nl) was introduced into a quartz capillary pre-filled with buffered electrolyte. After a high voltage (up to 30 kV) was applied to the ends of the capillary, the components of the mixture started to move at different speeds depending primarily on charge and mass (more precisely, the ionic radius) and accordingly reached the detection zone at different times. The following conditions were used: the total length of the capillary was 75 cm; the effective length (i.e. the length from the entrance to the detector window) was 65 cm; the operating voltage applied to the electrodes was +13 kV; the internal diameter of the capillary was 50 μm; detection was at 254 nm; temperature 200°C; sample injected under 300 mbar pressure; composition of the working buffer = 5 mM tartaric acid, 2 mM 18-crown-6. Sample preparation consisted of sample hydrolysis followed by dilution with the buffer solution.

### 2.8. Interaction with human serum albumin: circular dichroism

To estimate changes in protein structure upon addition of *R. canina* extracts, CD spectra were obtained. CD spectra from 1 µg ml^-1^ human serum albumin (HSA) were checked alone and with increasing concentrations of leaf and twig extracts. Measurements were made over the 260-195 nm wavelength range using a 0.5 cm path length Helma quartz cell. The recording parameters were as follows: scan speed, 50 nm min^-1^; step resolution, 0.5 nm; response time, 4 s; bandwidth, 1 nm; slit, auto. The CD spectra were corrected against a baseline with buffer only. The mean residue ellipticity θ (cm^2^ dmol^-1^) was calculated using software provided by Jasco.

### 2.9. Interaction with biological membranes of human cells; hemolysis test

The potential to damage cell membranes was investigated using the hemolysis method. Blood was collected from a Blood Bank in Lodz and centrifuged several times with phosphate buffered saline (PBS), pH 7.4, at 4°C. After washing, the hematocrit was measured and the blood samples were diluted to 2% haematocrit. The leaf and twig extracts of *Rosa canina* were added at concentrations of 0.5–50 µg/ml and left for 24 h at 37°C. After this incubation, hemolysis was measured at λ=540 nm using a BioTek plate reader and the values were calculated as follows:

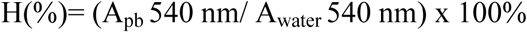

where A_pb_ is the absorbance of a tested sample, A_water_ means 100% hemolysis (erythrocytes incubated in distilled water only). Three independent measurements were obtained. The results are presented as mean ± SD.

### 2.10. Human plasma

Fresh human plasma was obtained from healthy, non-smoking volunteers. The blood was collected in tubes with CPD (citrate/phosphate/dextrose; 9:1 v/v blood/CPD). The fresh plasma was incubated (30 min, 37°C) with 0.5-50 μg ml^-1^ leaf and twig extracts and 4.7 mM H_2_O_2_/3.8 mM FeSO_4_/2.5 mM EDTA. Protein concentration was calculated from the absorbance at λ=280 nm using the Kalckar formula according to Whitaker and Granum (1980).

### 2.11. Lipid peroxidation

Lipid peroxidation products were determined with thiobarbituric acid (TBA) by measuring the TBARS concentration. After 30 min incubation of the samples (plasma plant extract H_2_O_2_/Fe) at 37°C, TCA and TBA were added and the mixtures were heated at 100°C, cooled and centrifuged. The absorbance of the supernatant at λ=535 nm was measured (SPECTROstar Nano Microplate Reader, BMG LABTECH, Germany). The TBARS concentration was calculated using the molar absorption coefficient (ε=156000 M^-1^ cm^-1^).

### 2.12. Carbonyl groups

Some ammonia derivatives bond covalently to carbonyl groups, so the product of oxidation of amino acid residues in the plasma proteins was carbonyl-depleted. This was used to assess oxidative damage.

2,4-dinitro-phenylhydrazine was used to identify carbonyl groups. After 1 h incubation in the dark at room temperature, dinitrophenylhydrazone (DNP) was formed. Its concentration was determined spectrophotometrically at λ=375 nm (SPECTROstar Nano Microplate Reader, BMG LABTECH, Germany). The carbonyl group concentration was calculated using the molar absorption coefficient (ε=22000 M^-1^ cm^-1^).

### 2.13. Thiol groups

The concentration of thiol groups was determined spectrophotometrically with Ellman reagent at λ=412 nm (SPECTROstar Nano Microplate Reader, BMG LABTECH, Germany) using the molar absorption coefficient (ε=13600 M^-1^ cm^-1^). The results were presented as nmol thiol groups mg^-1^ plasma protein.

### 2.14. Free radical scavenging

The free radical scavenging activity of leaf and twig extracts from *R. canina* was measured using the DPPH radical (2.2’-diphenyl-1-picrylhydrazyl, Sigma Aldrich). DPPH was dissolved in ethanol to a final concentration of 8.3×10^−5^ M. The antioxidant properties were tested at extract concentrations of 0.5–50 µg ml^-1^ and over different incubation times (after 5, 10, 15, 30 and 45 min). Absorbances were recorded at λ= 517 nm. Three independent repetitions were performed. The results are presented as percentage DPPH inhibition, calculated as:

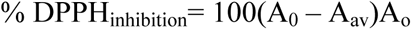

where A_0_ is the absorbance of DPPH solution and A_av_ is the average absorbance of samples treated with the extracts. The results are presented as means ± SD.

### 2.15. Cells

HEK 293 (normal human kidney) and BJ (normal human fibroblast) cell lines were purchased from ATCC (UK). The cells were cultured in DMEM (Gibco) supplemented with 10% fetal bovine serum and 1% streptomycin/penicillin. They were kept at 37°C in a humidified atmosphere.

### 2.16. Cytotoxicity

The cytotoxicity of the plant extracts was tested using the HEK293 cell line. The metabolic activity of the cells was checked using the Alamar Blue assay. The cells were seeded at 10,000 per well and left overnight to adhere. Next, 0.5–50 µg ml^-1^ *R. canina* leaf and twig extracts were added. After 24 h, incubation cytotoxicity was measured and viability was calculated as follows:

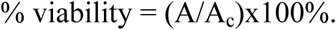

Where A = absorbance of a tested sample, A_c_ = absorbance of the control sample. Three independent measurements were collected and the results are presented as means ± SD.

### 2.17. ROS inhibition in human BJ cell line

The ability to decrease cellular ROS levels was tested using BJ cells. The cells were seeded at 2×10^5^ density and left for 24 h to adhere, then treated with 50 µg ml^-1^ leaf or twig extract. After 24 h incubation the medium was refreshed and 80 µM H_2_O_2_ was added for 30 min. After this incubation the cells were washed and 5 µM H_2_DCFDA was added to each sample for 20 min. The cells were washed with PBS and observed with a confocal microscope (Leica TCS SP8).

### 2.18. Statistical analysis

GraphPad Prism 5.0 and Statistica 13.1 were used for statistical analyses. Non-parametric ANOVA (Kruskal-Wallis test) was used to estimate the significance of differences. Significance was accepted when p < 0.05.

## 3. Results

### 3.1. HPLC analysis

HPLC analysis identified and quantified 40 phenolic compounds in the dried leaf and twig extracts (Table 1). Monomeric and polymeric catechins and gallic acid esters such as epigallocatechin were found in both extracts. There was more of the monomeric (+)-catechin content in twig extracts, whereas polymeric procyanidin B2 dominated in leaf extracts. There were high contents of dihydroxybenzoic and protocatechuic acids in the twig extracts. Both extracts had high ellagic acid contents, but leaves contained about twice as much as twigs. From the flavonoid group, leaf extracts had more cyanidin than twig extracts, and also contained high levels of neochlorogenic acid.

**Table.1.** Content of phenolic compounds detected using HPLC in dried extracts of *R. canina* leaves and twigs.

| No. | Phenolic compound | Content, [mg/g] in dry matter of extract |  |
| --- | --- | --- | --- |
|  |  | Leaf | Twig |
| 1 | Gallic acid (3.4.5-Trihydroxybenzoic acid) | 0.805 | 0.357 |
| 2 | *p*-Benzoquinone (Quinone) | 0.252 | 0.989 |
| 3 | α-resorcylic acid (3.5-Dihydroxybenzoic acid) | 0.138 | 0.117 |
| 4 | Pyrocatechol (1.2-Dihydroxybenzene. Catechol) | 0.226 | 0.493 |
| 5 | Protocatechuic acid (3.4-Dihydroxybenzoic acid) | 0.153 | 13.911 |
| 6 | Neochlorogenic acid (*trans*-5-*O*-Caffeoylquinic acid) | 57.148 | 0.258 |
| 7 | (-)-epigallocatechin (monomeric flavan-3-ol) | 0.207 | 1.680 |
| 8 | (+)-Catechin (flavan-3-ol) (monomeric flavan-3-ol) | 2.804 | 17.798 |
| 9 | 4-hydroxybenzoic acid | 1.182 | 0.323 |
| 10 | Procyanidin B2 (pentahydroxyflavane); (*cis.cis*"-4.8"-Bi(3.3'.4'.5.7-pentahydroxyflavane) (Polymeric flavan-3-ol) | 22.473 | 3.222 |
| 11 | Gentisic acid (2.5-Dihydroxybenzoic acid) | 1.577 | 0.340 |
| 12 | 4-Hydroxybenzaldehyde | 1.111 | 0.263 |
| 13 | Chlorogenic acid (*trans*-3-*O*-Caffeoylquinic acid) | 4.609 | 0.934 |
| 14 | Vanilic acid (4-Hydroxy-3-methoxybenzoic acid) | 0.102 | 0.379 |
| 15 | β-resorcylic acid (2.4-Dihydroxybenzoic acid) | 0.017 | 0.015 |
| 16 | Caffeic acid (*trans*-3.4-Dihydroxycinnamic acid) | 0.035 | 0.203 |
| 17 | (-)-epicatechin ((-)-*cis*-3.3'.4'.5.7-Pentahydroxyflavane) (monomeric flavan-3-ol) | 1.822 | 1.379 |
| 18 | Syringic acid (4-Hydroxy-3.5-dimethoxybenzoic acid) | 0.613 | 0.134 |
| 19 | 1.3-Dicaffeoylquinic acid | 0.826 | 0.189 |
| 20 | Cyanidin (3.3'.4.5.7-Pentahydroxyflavone) (3.3'.4.5.7-Pentahydroxyflavylium chloride) | 47.448 | 4.453 |
| 21 | Syringaldehyde (4-Hydroxy-3.5-dimethoxybenzaldehyde) | 0.402 | 0.262 |
| 22 | *p*-Coumaric acid (*trans*-4-Hydroxycinnamic acid) | 0.520 | 0.247 |
| 23 | Ferulic acid (4-Hydroxy-3-methoxy-cinnamic acid) | 0.439 | 0.081 |
| 24 | Coumarin (1.2-Benzopyrone) | 1.285 | 0.170 |
| 25 | Sinapic acid (4-Hydroxy-3.5-dimethoxy-cinnamic acid) | 0.214 | 0.140 |
| 26 | *trans*-3-Hydroxycinnamic acid (*m*-Coumaric acid) | 0.241 | 0.119 |
| 27 | Luteolin 7-*O*- β-*D*-glucoside | 1.614 | 1.417 |
| 28 | Rutin (quercetin-3-*O*-rutinoside) | 26.661 | 4.431 |
| 29 | Ellagic acid (4.4'.5.5'.6.6'-Hexahydroxydiphenic acid) | 35.881 | 14.448 |
|  | 2.6.2'.6'-dilactone) |  |  |
| 30 | Hesperidin (Hesperetin-7-rutinoside) | 4.013 | 0.633 |
| 32 | *o*-Coumaric acid (*trans*-2-Hydroxycinnamic acid) | 0.711 | 0.149 |
| 31 | Rosmarinic acid 3.4-Dihydroxycinnamic acid (*R*)-1-carboxy-2-(3.4-dihydroxyphenyl)ethyl ester | 4.406 | 1.851 |
| 33 | Salicylic acid (2-Hydroxybenzoic acid) | 0.457 | 0.474 |
| 34 | Myricetin (flavonol) (3.3'.4'.5.5'.7-Hexahydroxyflavone) | 3.928 | 7.175 |
| 35 | Quercetin (flavonol) (3.3'.4'.5.7-Pentahydroxyflavone) | 0.156 | 0.241 |
| 36 | *trans*-Cinnamic acid | 0.167 | 0.034 |
| 37 | Naringenin (4'.5.7-Trihydroxyflavanone) | 0.461 | 0.924 |
| 38 | Luteolin (3'.4'.5.7-Tetrahydroxyflavone) | 0.911 | 0.364 |
| 39 | Kaempferol (3.4'.5.7-Tetrahydroxyflavone) | 0.148 | 0.167 |
| 40 | 3-Hydroxyflavone | 0.103 | 0.128 |

### 3.2. Phenolic compounds

There was no difference in total phenolic compound contents between the water extracts of *R. canina* leaves and twigs (Fig. 1). Flavonoids accounted for about 9.5% and 5.5% of the phenolic compounds, respectively. However, because AlCl_3_ reacts mainly with flavones, flavonols, flavanones and flavanonols, the result does not correspond exactly to the total flavonoid content of the extracts. The total catechin (flavan-3-ol) content of the twig extracts was about twice that of the leaf extracts. Among the phenolic compounds, catechins constituted about 5.2% and 10% in leaf and twig extracts, respectively.

**Figure 1.**
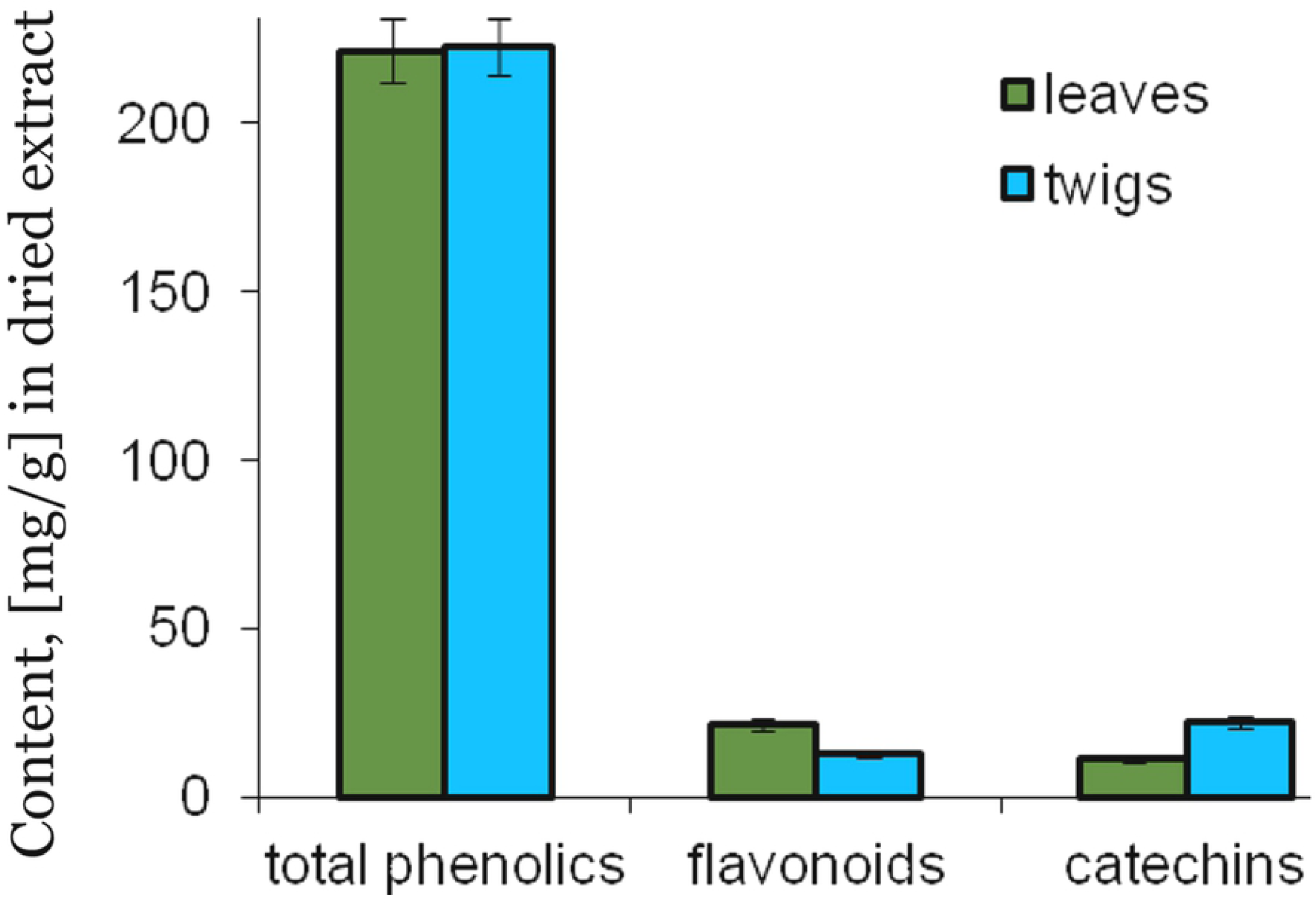
Phenolic compound content of the extracts of *R. canina* leaves and twigs. Values are means ± SD (n=3).

### 3.3. Water-soluble vitamins

Table 2 shows the contents of five B vitamins in the leaves and twigs of *R. canina*. In all tested samples the predominant B vitamin was B6 (pyridoxine). The amounts of each vitamin were similar in both organs of *R. canina* investigated.

**Table 2.** Total content of water soluble vitamins in dry matter of the extracts from *R. canina* twigs and leaves.

| Vitamins | Content, [%] in dry matter of the extracts |  |
| --- | --- | --- |
|  | Leaf | Twig |
| B1 (thiaminechloride) | 0.112 | 0.077 |
| B2 (Riboflavin) | 0.048 | 0.051 |
| B3 (Pantothenicacid) | 0.21 | 0.27 |
| B5 (nicotinicacid) | 0.33 | 0.31 |
| B6 (pyridoxine) | 0.57 | 0.62 |
| Bc (folicacid) | 0.097 | 0.086 |

### 3.4. Determination of vitamin E content

More tocopherols were found in the leaf than twig extract of *R. canina*. The concentration of isomer α was highest in both parts of the plants (Table 3).

**Table 3.**
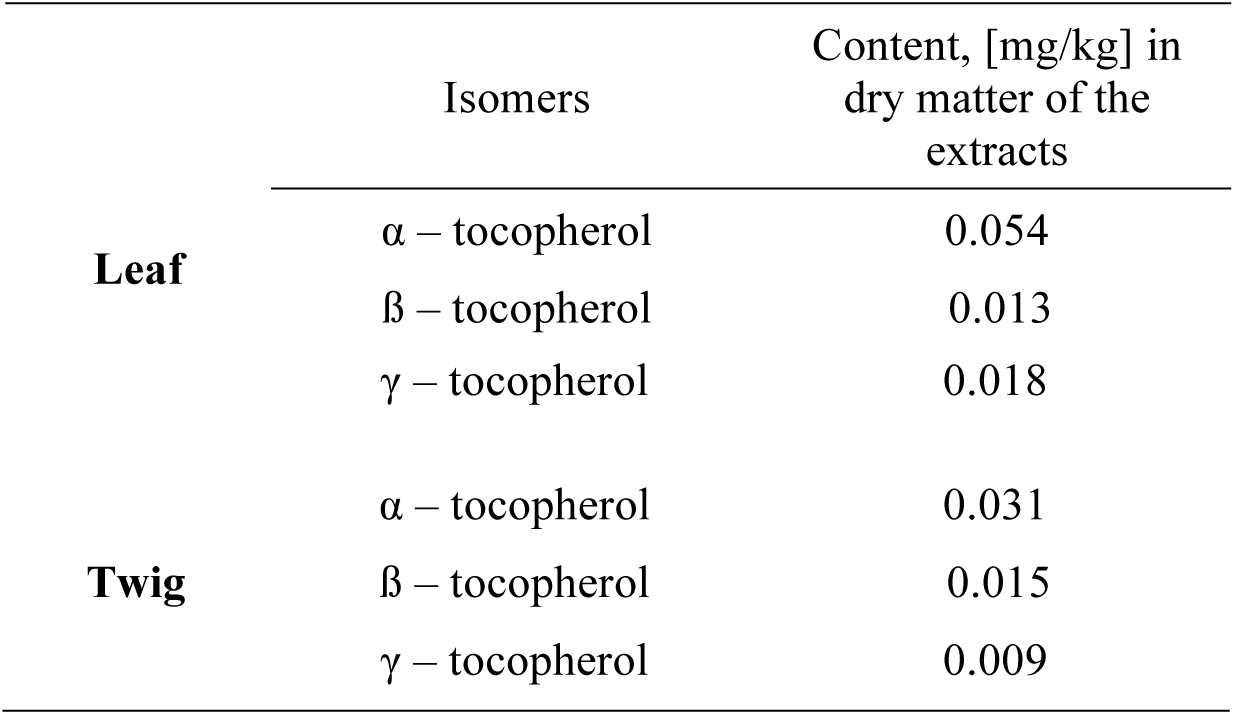
Total content of vitamin E isomers in the leaf and twig extracts from *R. canina*.

### 3.5. Amino acids content

On the basis of the values presented in Table 4, 13 amino acids were identified in the leaves and twigs of *R. canina*, nine of them essential: valine, threonine, methionine, isoleucine, leucine, phenylalanine, histidine, lysine and arginine. The total content of amino acids was higher in leaves (6.1487%) than twigs (1.6864%). The highest concentrations were of proline (0.3196%), serine (0.2040%) and phenylalanine (0.2262%) in the twig extracts, and of proline (1.04994%), valine (0.6046%) and phenylalanine (0.6045%) in the leaf extracts.

**Table 4.**
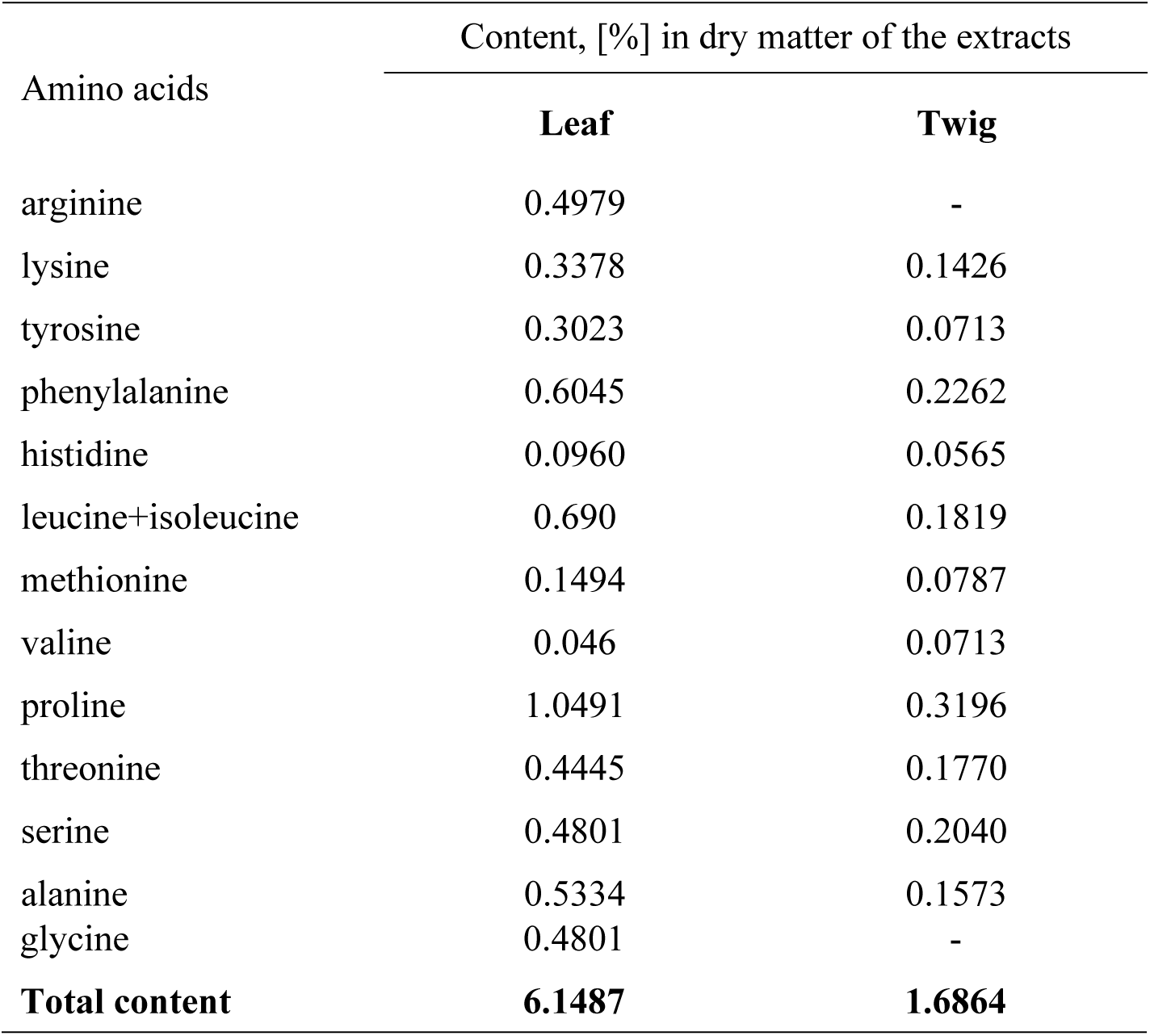
The content of amino acids (%) in dried extracts from twigs and leaves of *R. canina*

| Amino acids | Content, [%] in dry matter of the extracts |  |
| --- | --- | --- |
|  | Leaf | Twig |
| arginine | 0.4979 | - |
| lysine | 0.3378 | 0.1426 |
| tyrosine | 0.3023 | 0.0713 |
| phenylalanine | 0.6045 | 0.2262 |
| histidine | 0.0960 | 0.0565 |
| leucine+isoleucine | 0.690 | 0.1819 |
| methionine | 0.1494 | 0.0787 |
| valine | 0.046 | 0.0713 |
| proline | 1.0491 | 0.3196 |
| threonine | 0.4445 | 0.1770 |
| serine | 0.4801 | 0.2040 |
| alanine | 0.5334 | 0.1573 |
| glycine | 0.4801 | - |
| **Total content** | **6.1487** | **1.6864** |

### 3.6. Interaction with human serum albumin: Circular dichroism

Changes in the secondary structure of HSA in the presence of *R. canina* leaf and twig extracts were checked using circular dichroism. CD spectra for HSA at pH 7.4 were obtained in the absence and presence of the extracts.

The HSA CD spectrum contained two characteristic negative bands in the far UV at 202 and 220 nm (Fig. 2). As increasing concentrations of the extracts were added the amount of α-helix decreased whereas the amounts of β-sheet and random coil increased. The leaf extract changed the albumin structure more markedly than the twig. Percentage values calculated using CDNN software indicated a decrease in α-helical structure from 62.1% to 21.4% for the leaf extract and from 58.9% to 40.7% for the twig extract. The amount of β-sheet increased from 12.6% to 19.4%, and that of random coil from 15.8% to 38.5% and from 17.2% to 24.8% in the leaf and twig extracts, respectively (Table 5).

**Table 5.**
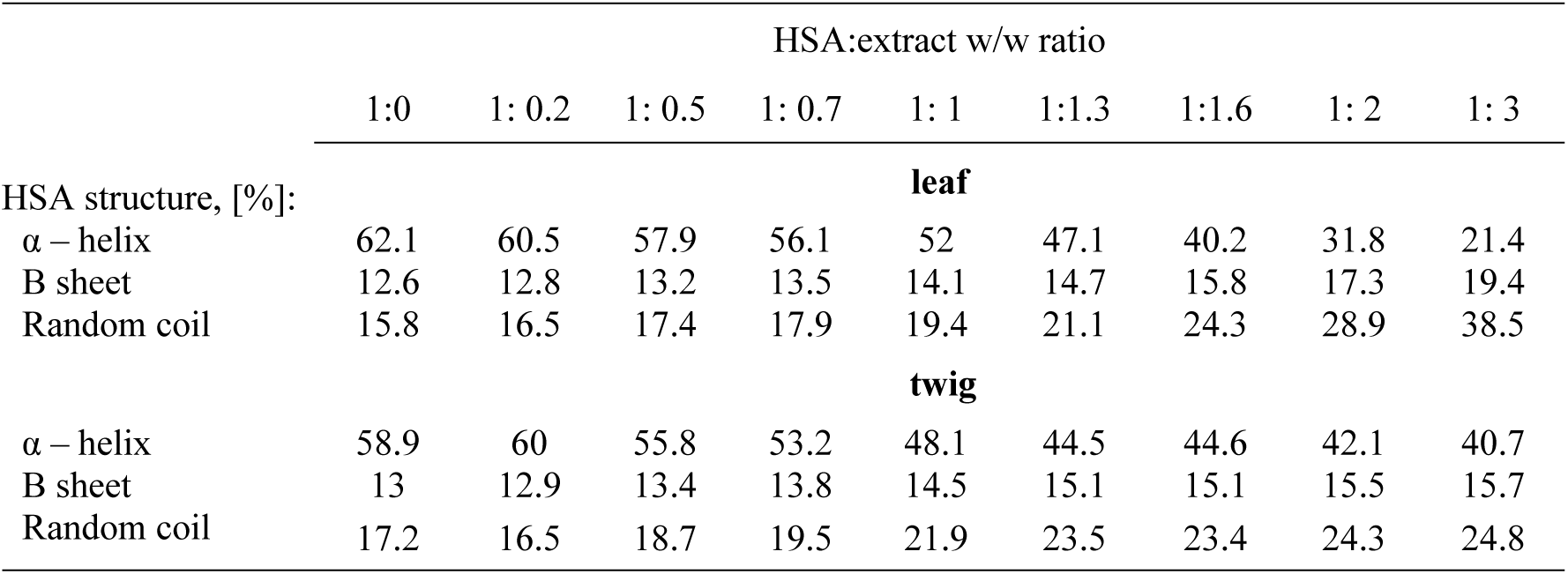
The changes in secondary structure of HSA in the presence of different ratios (w/w) of HSA:*R. canina* extracts.

|  | HSA:extract w/w ratio |  |  |  |  |  |  |  |  |
| --- | --- | --- | --- | --- | --- | --- | --- | --- | --- |
|  | 1:0 | 1: 0.2 | 1: 0.5 | 1: 0.7 | 1: 1 | 1:1.3 | 1:1.6 | 1: 2 | 1: 3 |
| HSA structure, [%]: |  |  |  |  |  |  |  |  |  |
|  |  |  |  |  | **leaf** |  |  |  |  |
| $\alpha$ – helix | 62.1 | 60.5 | 57.9 | 56.1 | 52 | 47.1 | 40.2 | 31.8 | 21.4 |
| B sheet | 12.6 | 12.8 | 13.2 | 13.5 | 14.1 | 14.7 | 15.8 | 17.3 | 19.4 |
| Random coil | 15.8 | 16.5 | 17.4 | 17.9 | 19.4 | 21.1 | 24.3 | 28.9 | 38.5 |
|  |  |  |  |  | **twig** |  |  |  |  |
| $\alpha$ – helix | 58.9 | 60 | 55.8 | 53.2 | 48.1 | 44.5 | 44.6 | 42.1 | 40.7 |
| B sheet | 13 | 12.9 | 13.4 | 13.8 | 14.5 | 15.1 | 15.1 | 15.5 | 15.7 |
| Random coil | 17.2 | 16.5 | 18.7 | 19.5 | 21.9 | 23.5 | 23.4 | 24.3 | 24.8 |

**Figure 2.**
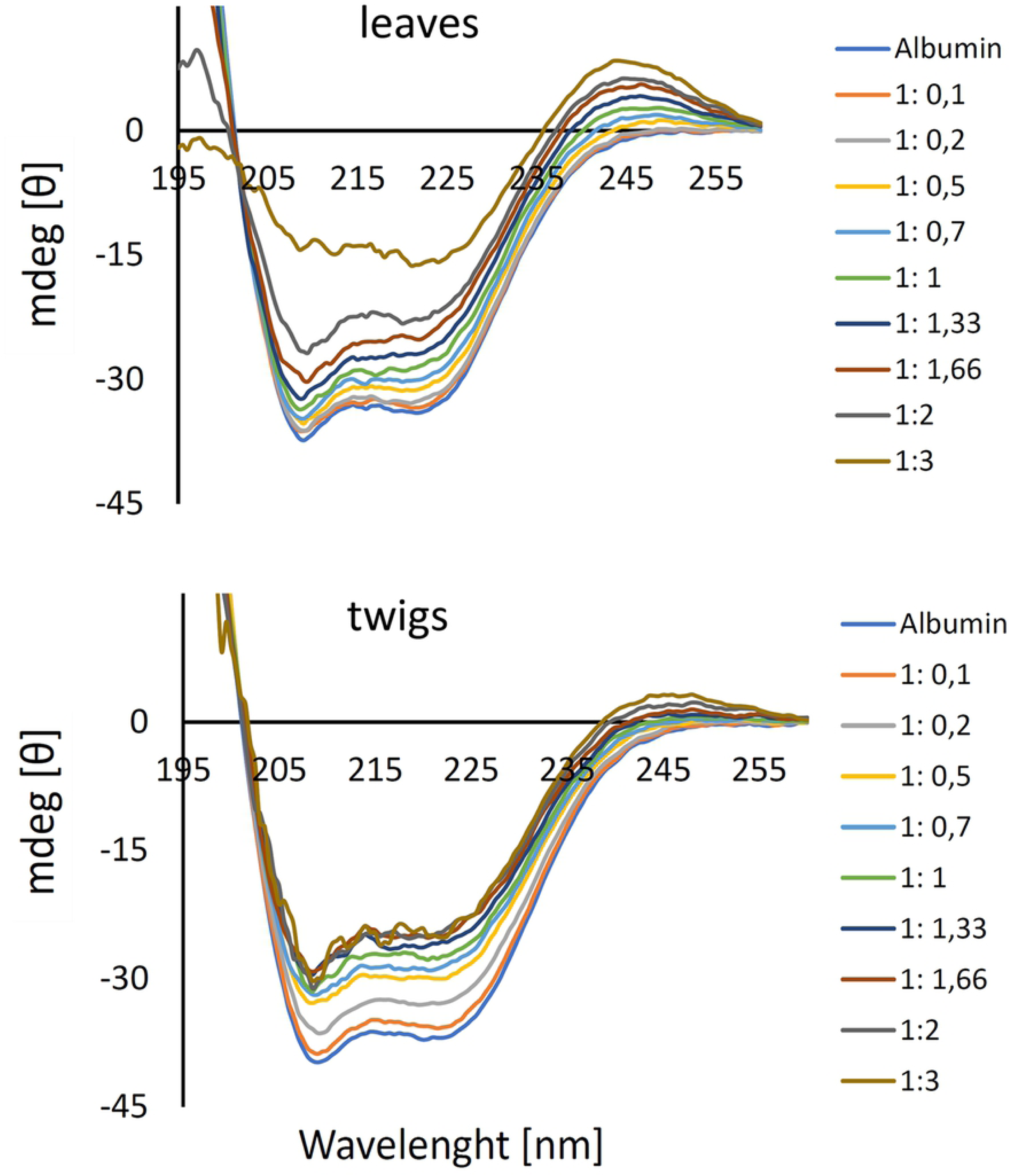
Ellipticity changes of human serum albumin (1 µM) in the presence of varying ratios (1:0.1 – 1:3) of *R. canina* leaf and twig extracts.

### 3.7. Interaction with erythrocyte membranes; hematoxicity

The *R. canina* leaf and twig extracts were subjected to a hemolysis test. Both proved non-toxic for erythrocytes, and the hemotoxicity did not exceed 5% even at the highest extract concentrations (Fig. 3).

**Figure 3.**
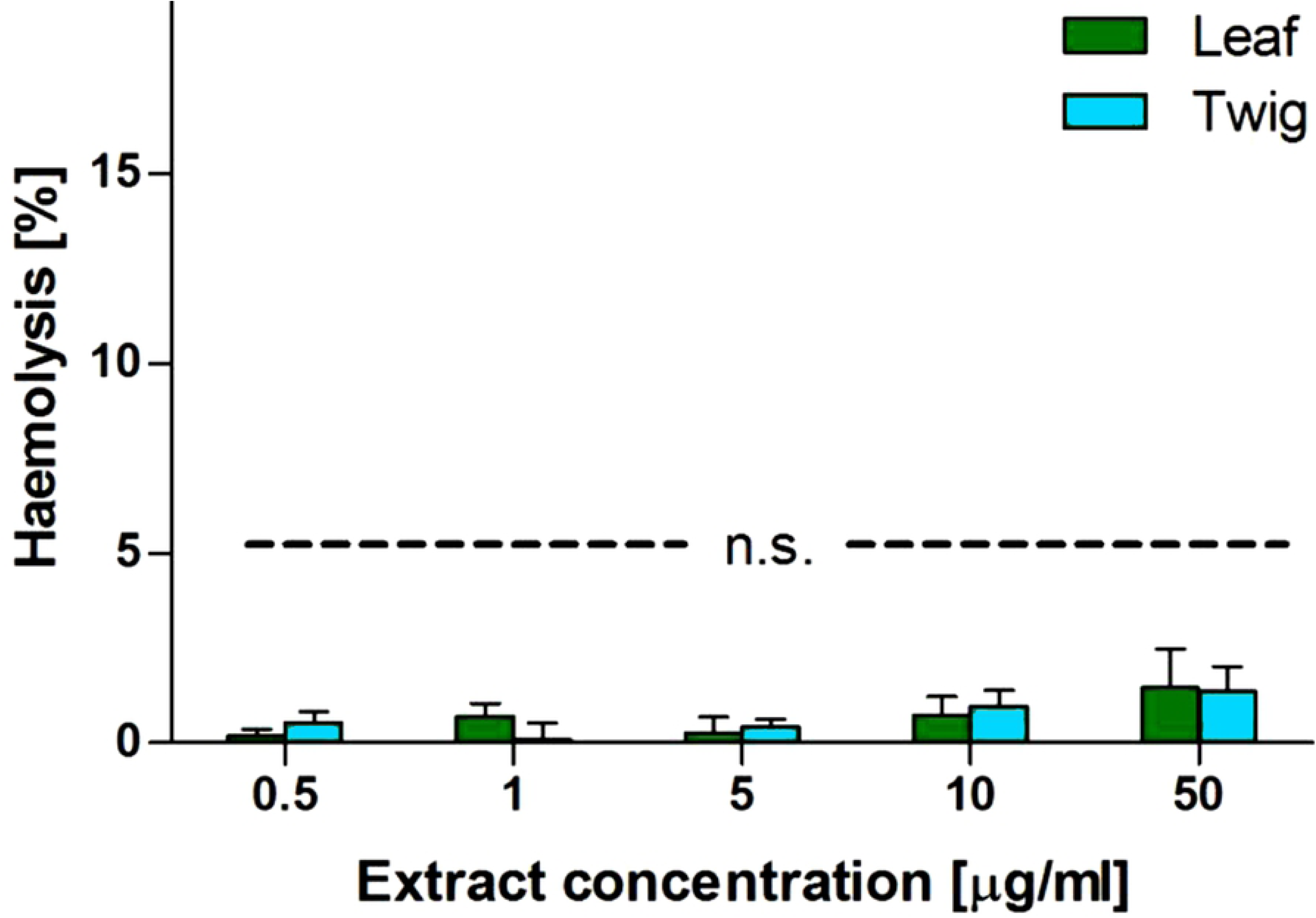
Hemolysis after 24 h incubation with the *R. canina* leaf and twig extracts. Data are mean ± SD from three independent measurements. Ns – not statistically significant.

### 3.8. Cytotoxicity

The cytotoxicity of the extracts was tested on the HEK 293 cell line (Fig. 4). The results indicate that the extracts were non-toxic toward these cells. There were no significant differences between the leaf and twig samples.

**Figure 4.**
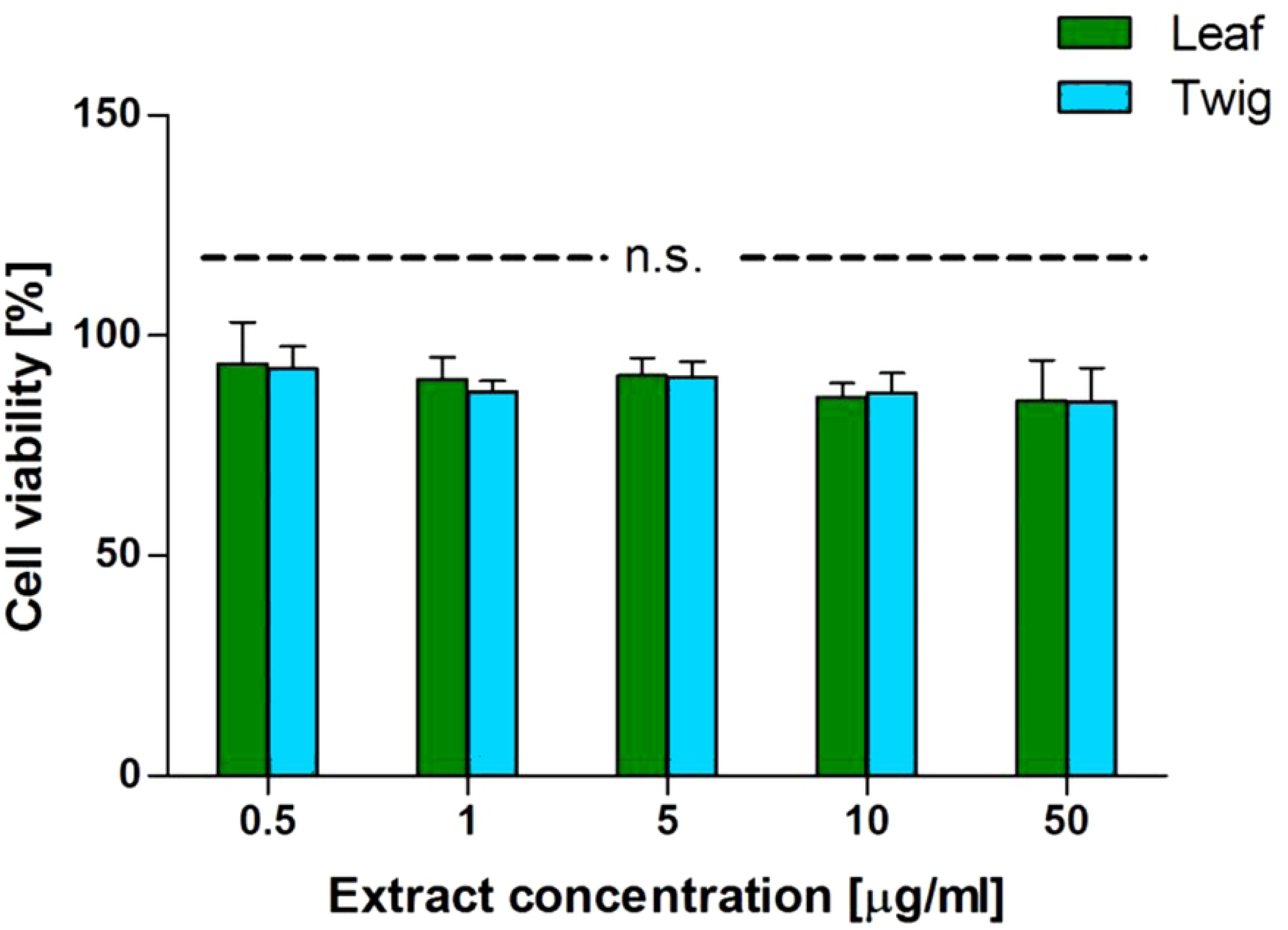
HEK 293 cell line viability after 24 h incubation with *R. canina* leaf and twig extracts. Data are mean ± SD from three independent measurements. Ns – not statistically significant.

### 3.9. Markers of oxidative stress

The tested extracts caused no statistically significant change in plasma lipid peroxidation (Fig. 5), nor did they affect thiol groups in plasma proteins treated with H_2_O_2_/Fe (Fig. 6). Fig. 7 shows that the leaf extract at the highest concentration and the twig extract at 10 and 50 μg ml^-1^ inhibited H_2_O_2_/Fe-induced protein carbonylation.

**Figure 5.**
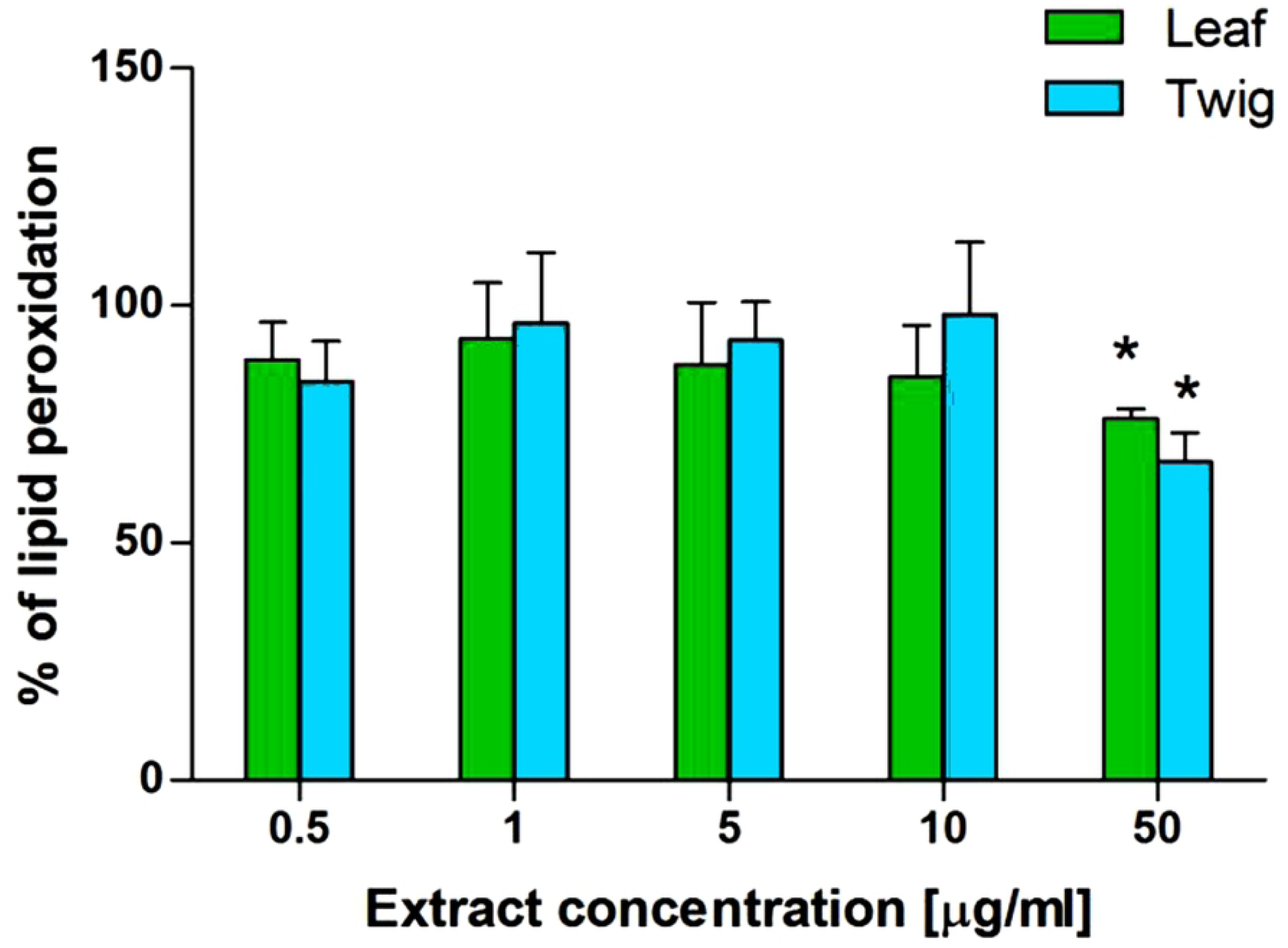
Effect of the *R. canina* twig and leaf extracts (0.5–50 μg/ml) on H_2_O_2_/Fe-induced plasma lipid peroxidation. The data are mean ± SEM of three independent repeats. Ns – not statistically significant.

**Figure 6.**
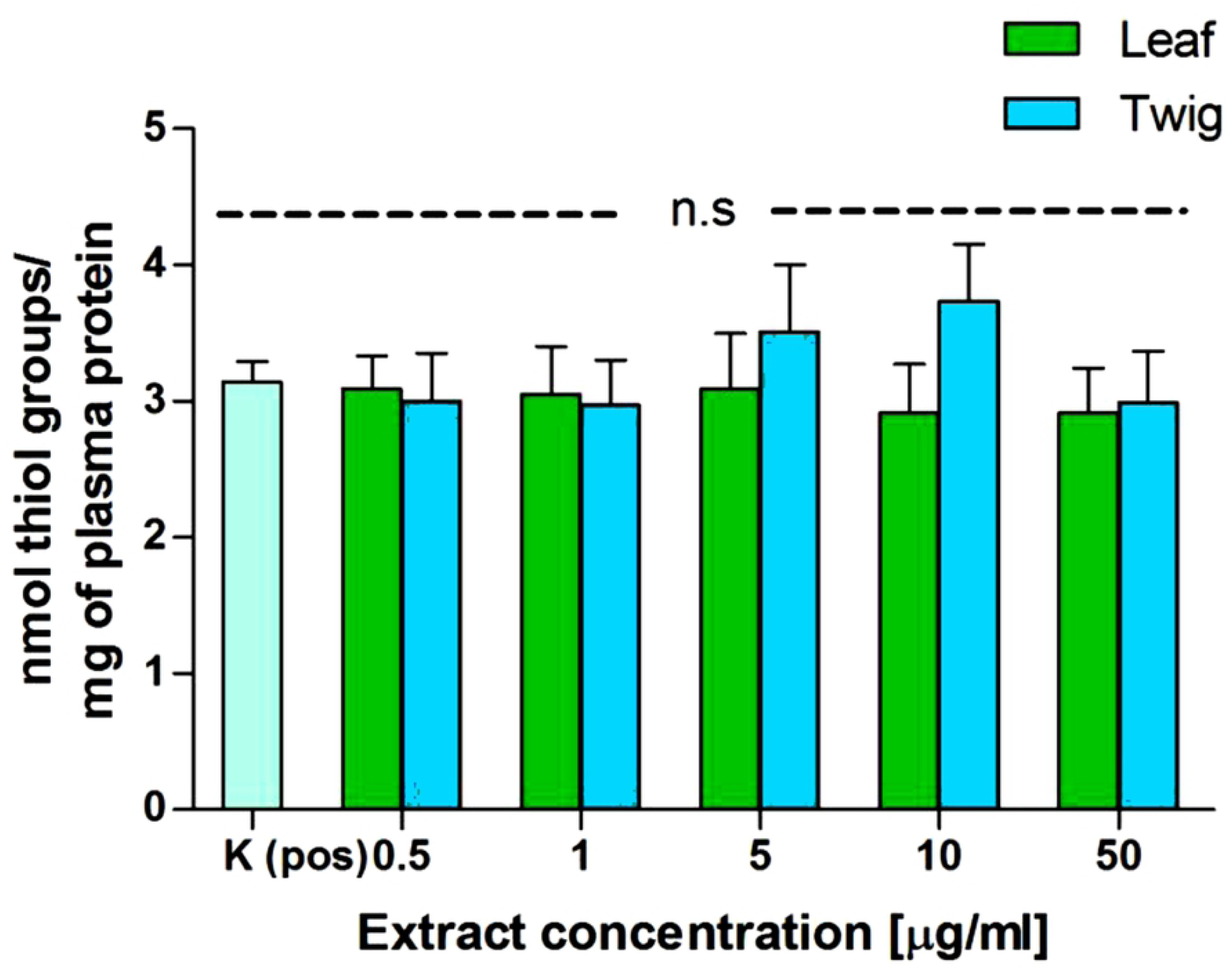
Effect of the *R. canina* twig and leaf extracts (0.5–50 μg/ml) on H_2_O_2_/Fe-induced oxidation of thiol groups. The data are mean ± SEM of three independent repeats. Ns – not statistically significant.

**Figure 7.**
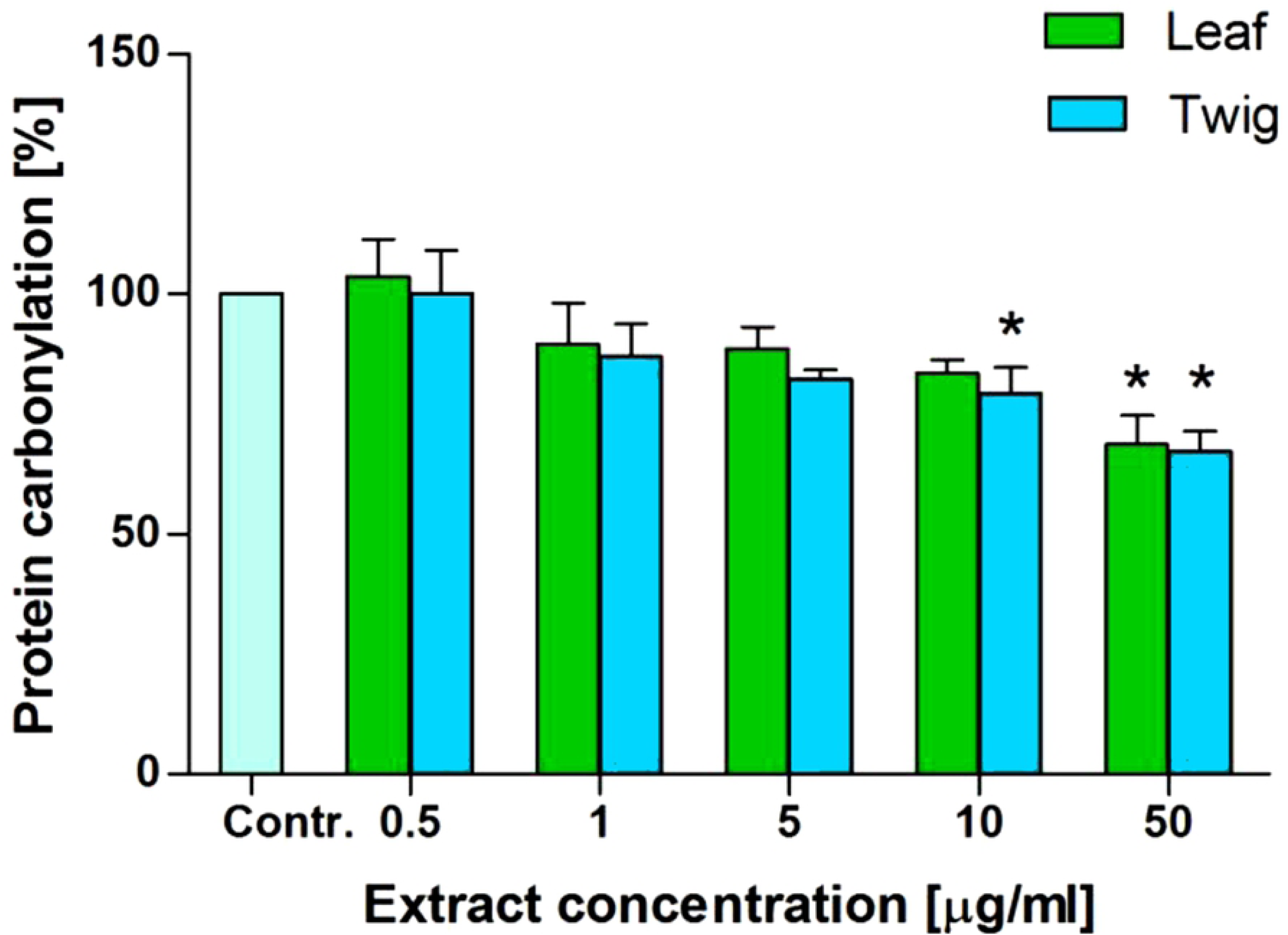
Effect of *R. canina* twig and leaf extracts (0.5–50 μg/ml) on protein carbonylation induced by H_2_O_2_/Fe. The data are means ± SEM of three independent repeats. * - p< 0.05 vs. control.

### 3.10. Free radical scavenging

The scavenging activity of the extracts was measured by the DPPH method. Both the leaf and twig extracts from *R. canina* exhibited antiradical activity. The differences between the activities of the extracts were biggest at 5 and 10 µg/ml concentrations (Table 6): the leaf extracts caused 26% and 41% inhibition while the twig extracts caused 13% and 36% inhibition.

**Table 6.** Percentages of free radical scavenging by 0.5–50 µg ml^-1^ *R. canina* leaf and twig extracts during 10, 15 and 45 min incubations. The results are means ± SD from three independent measurements.

| Incubation time | 0.5 µg/ml | 1 µg/ml | 5 µg/ml | 10 µg/ml | 50 µg/ml |
| --- | --- | --- | --- | --- | --- |
| **Leaf** |  |  |  |  |  |
| 10 min | 0.6 ± 0.2 | 0.8 ± 0.1 | 20.5 ± 0.5 | 32.9 ± 1.8 | 72.0 ± 1.5 |
| 15 min | 0.7 ± 0.4 | 0.7 ± 0.1 | 24.3 ± 0.4 | 42.2 ± 1.9 | 71.9 ± 2.0 |
| 45 min | 0.2 ± 0.4 | 0.0 ± 0.1 | 26.2 ± 0.6 | 41.9 ± 2.3 | 70.9 ± 2.1 |
| **Twig** |  |  |  |  |  |
| 10 min | 0.0 ± 0.3 | 0.5 ± 0.1 | 12.6 ± 0.5 | 34 ± 1.8 | 72.8 ± 1.5 |
| 15 min | 0.1 ± 0.4 | 0.9 ± 0.1 | 11 ± 0.4 | 33.7 ± 1.9 | 71.7 ± 2.1 |
| 45 min | 0.7 ± 0.4 | 0.5 ± 0.1 | 13 ± 0.6 | 36.8 ± 2.3 | 71.3 ± 2.2 |

### 3.11. ROS inhibition

The ability of *R. canina* leaf and twig extracts to decrease ROS production in human fibroblasts was tested using a H_2_DCFDA probe. Both extracts protected the cells by inhibiting ROS production, though the leaf extract had the stronger effect of the two (Fig. 8).

**Figure 8.**
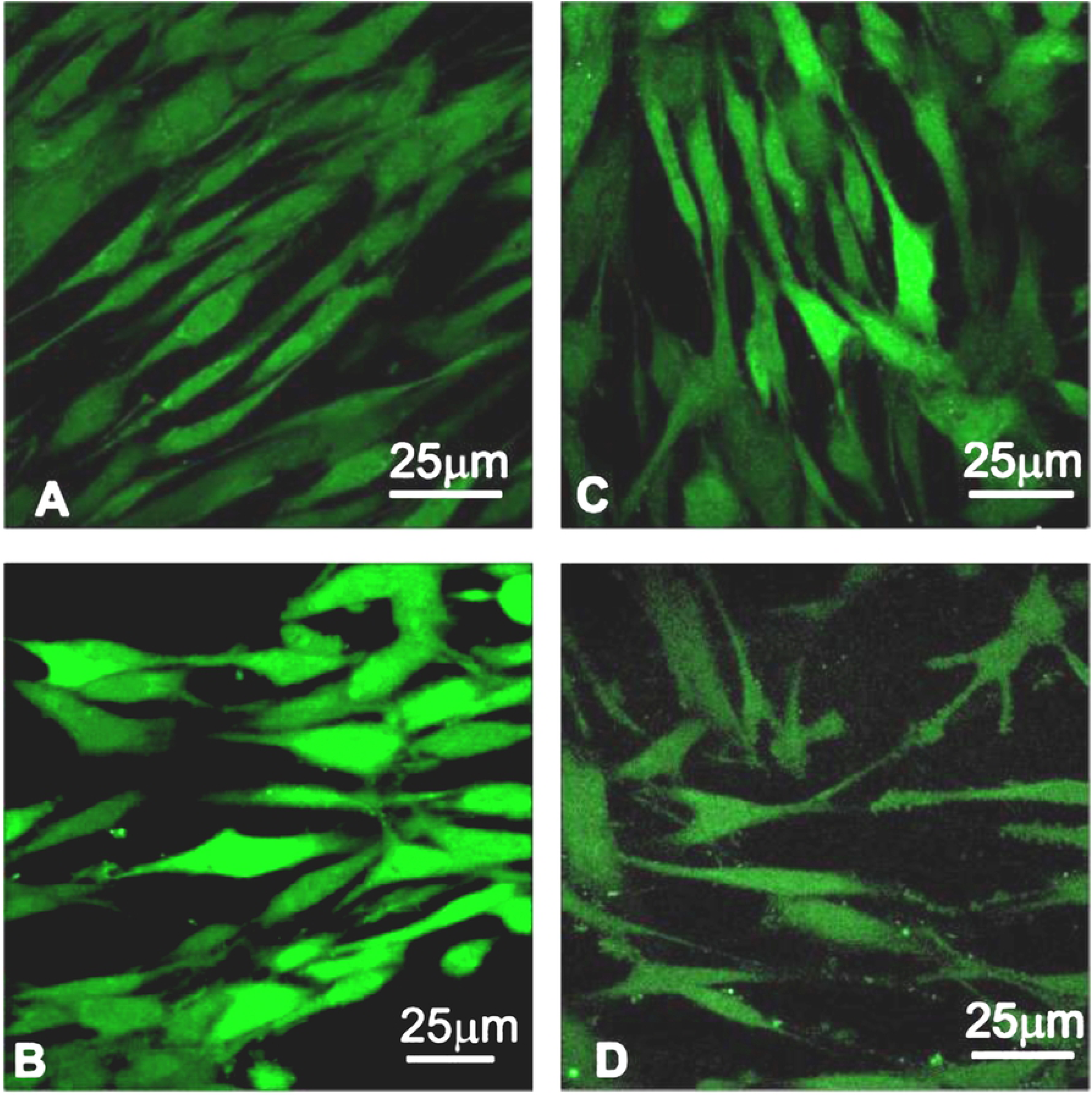
Confocal microscopy images of BJ cells after 24 h treatment with the *R. canina* leaf and twig extracts. (A) – control; (B) – H_2_O_2_ 80 µM; (C) – H_2_O_2_ + twig extract, 50 µg ml^-1^; (D) - H_2_O_2_ + leaf extract, 50 µM. Scale bar=25 µm.

## 4. Discussion

Many reports describe *Rosa canina* L. hips as an abundant source of antioxidants [1–4]. Here we demonstrated that other parts of this plant, the twigs and leaves, can be considered potential sources of compounds with beneficial properties for humans, including antioxidant and antiradical activities.

The difference in the phenolic compound profile between *Rosa sp*. twigs and leaves is well documented [5]. The twigs are a very rich source of catechins, cyanidin derivatives, and also contain quercetin and quinic acid derivatives, while the leaves are rich in quercetin rhamnoside, catechins and proanthocyanidin secondary metabolites. Among the *Rosaceae* plants examined, the highest amounts of catechins and phenolic acids were found in *R. canina* leaves [5].

HPLC analysis revealed over 40 different phenolic compounds in the *R. canina* leaves and twigs. The leaves used in these studies were rich in chlorogenic and neochlorogenic acid, cyanidin and procyanidin B2, ellagic acid, rutin (quercetin-3-O-rutinoside), rosmarinic acid, myricetin, epicatechin and coumarin, while the twigs had relatively high levels of ellagic acid, cyanidin, myricetin and rutin and were rich in protocatechuic acid, catechin and rosmarinic acid.

These results were consistent with those of Ouerghemmi et al., who showed that *R. canina* twig extracts contain catechin, rutin, quercetin and kaempferol, but we also found higher amounts of protocatechuic acid, cyanidin, ellagic acid and myricetin [37]. According to the latest studies, *Rosa sp*. twigs are a very rich source of catechins, cyanidin derivatives, and also contain quercetin and quinic derivatives and a very low amount of kaempferol, while leaves from this plant are rich in quercetin rhamnoside [5]. Other analyses of extracts from different *Rosa* species revealed that leaf extracts are rich in catechins and proanthocyanidins. Among the different *Rosaceae* plants examined, *R. canina* leaves contained the highest amounts of catechins and phenolic acids.

The quantity and quality profiles of phenolic compounds are likely to differ among *Rosa* species including *R. canina*, and among their organs [1]. Neochlorogenic acid is best-known phenolic acid, found in the many *Rosa* species extracts [1]. However, higher levels of this compound were found in *R. canina* leaves (57.148 mg g^-1^ DW) than twigs (0.2579 mg g^-1^ DW). Conversely, the *R. canina* twigs contained twice as much myricetin as the leaf extracts.

The phenolic compound profiles of the *R. canina* leaf and twigs extracts differed, and both differed from the profile described for hips [31]. Compared to the values of phenolic compounds in dried *R. canina* fruits reported by Kerasioti et al. [31], the leaf and twig extracts differed mainly in the quantities of selected phenolics. The levels of (+)-catechin in leaf and twig extracts were 48- and 7.57-fold lower, respectively, than in the fruit extracts. In contrast, the level of protocatechuic acid in twigs was 6.65-fold higher than reported for hips [31] and 10-fold higher than in leaves. The leaves contained a similar rutin concentration (26.661 mg g^-1^ DW) to hips (25.64 mg g^-1^ DW). Interestingly, chlorogenic acid, which is absent from dried *R. canina* fruits [31], is present in the extracts from dried leaves (4.6099 mg g^-1^ DW) and twigs (0.9339 mg g^-1^ DW).

The vitamin contents of *R. canina* leaves and twigs are still not well established. It is generally believed that the strongest contributor to the antioxidant properties of rose hips is the high vitamin C content. However, the hips from *R. canina* and *R. rugosa* are also very rich sources of tocopherols, which protect lipids against peroxidation [13]. Our study revealed tocopherol isoforms in both organs examined, though there was more in the leaves than the twigs. Also, only two isoforms were detected in hips, but all isoforms (α, β and γ tocopherols) were found in *R. canina* twigs and leaves in the present study.

The mean total tocopherol contents of hips were reported as 15.9 ± 1.7 μmol 100 g^-1^ in raw *R. canina*, 31.4 ± 3.2 μmol 100 g^-1^ in *R. canina* powder, and 8.7 ± 1.1 μmol 100 g^-1^ in *R. canina* puree. In the fleshy parts of ose hips only α- and γ–tocopherol were found, which indicates limited biosynthesis of δ-tocopherol and of tocotrienols during ripening. It is suggesed that the γ- and δ-tocopherols are converted by the action of γ-tocopherol methyltransferase to α- and β-tocopherol, respectively [42]. Although there is no information about the total tocopherol content in *R. canina* leaves and twigs, the present study showed the presence of tocopherol isoforms, more in the leaves than in the twigs.

B vitamins are not only very important in the human diet but are also considered as antioxidants in plants. Some experiments indicate that they accumulate mainly in seeds, but they can also be found in other parts of plants including leaves and twigs. The tested extracts exhibited relatively low concentrations of B vitamins, B6 (pyridoxine) being the most abundant.

Overall, since the leaves and twigs contain tocopherol isoforms and B vitamins, these parts of the plant can be considered as good sources of antioxidants [13,16,43]. Some amino acids also protect cells from oxidative stress, free radicals and heavy metals. Moreover, they mediate the synthesis of molecules such as glutathione, which is very important for the antioxidative response [44–46]. The total amino acid content of the extracts from *R. canina* leaves and twigs was measured in this study.

For non-meat consumers, the important information is that there were more amino acids in the leaf extracts than the twig extract (6.1 vs 1.6%, respectively). Both extracts contained essential amino acids such as lysine, phenylalanine, histidine, leucine, isoleucine, methionine, valine and threonine. Arginine was found only in the leaf extract and tryptophan was detected in neither. The total amino acid content was 61.48 mg g^-1^ DW in the rose leaf extract and 16.86 mg g^-1^ DW in the twig extract, so the leaves seem a good source of protein. According to WHO standards, *P. indica, P. hirta* and *E. thymifolia* could serve as good sources of protein. The total amino acid contents were 58.80 mg g^-1^ DW in *P. indica*, 123.92 mg g^-1^ DW in *E. thymifolia*, and 225.73 mg g^-1^ DW in *P. hirta* [14.15]. Although the leaf extract was richer in amino acids, the ratio of essential to total amino acids was 0.37, whereas in the twig extract it was 0.55.

Serum albumin transports many compounds in the blood. Thus, it was important to check interactions between albumin and the studied extracts. We examined the conformational changes in human serum albumin (HSA) during titration with the water extracts. Circular dichroism is useful for checking how compounds interact with protein secondary structures. The minima characteristic of α-helices were visible around 202 and 222 nm in HSA. With increasing concentrations of the extracts the ellipticity increased.

The extracts changed the α-helix content and increased the amounts other protein structures such as β helix and random coil. The results suggest that compounds in the extracts can interact with proteins and cause conformational changes. The leaf extract interacted more strongly with HSA than the twig extract. Almost twice as much α-helix was lost when the leaf extract was used (final content of α-helix was 21.4% for leaf and 40.7% for twig extract). Das at al. reported that extracts containing flavonoids (quercetin, myricetin, kaempferol) could interact with HSA. Slight changes in albumin structure were observed when a 1:20 ratio was used [47]. The changes were more obvious when the maximum albumin:extract ratio was 1:3.

Plants can be considered good sources of antiradical agents and can be included in human diets. The antioxidant potential of the water extracts from *R. canina* leaves and twigs was demonstrated by several methods. While the antioxidant properties of rose hips are well documented, there are few reports about such properties in leaf and twig extracts [1,2,7,16]. We found that the extracts decreased the lipid peroxidation level slightly when the highest concentration was used, though this was non-significant; the extracts did not affect thiol group oxidation. However, both extracts at higher concentrations reduced protein carbonylation significantly.

DPPH and the ROS inhibition assay were used to confirm antiradical and antioxidant activities in the *R. canina* extracts. Both methods revealed that the leaf and twig extracts were highly effective. The free radical activity of DPPH was reduced when 5 µg ml^-1^ extracts were used. The data show that the extracts almost immediately inhibited DPPH activity and the free radical scavenging effect was most pronounced at 50 µg ml^-1^ concentration. Similar results were obtained when ROS inhibition was measured, but the leaf extract protected fibroblasts more effectively than the twig extracts. According to the literature, leaf extract from *R. canina* has one of the highest antioxidant potentials among the *Rosaceae* family, perhaps because *R. canina* has the highest phenolic content of all *Rosaceae*. Moreover, although the results presented in this paper were obtained with water extracts not methanol extracts as in other studies, all these results indicate that extracts from *R. canina* could have valuable antioxidant properties.

Antioxidant activity has not been so extensively investigated in twig extracts. Nevertheless, Ouerghemmi et al. [37] tested twig extracts from *R. canina, R. sempervirens* and *R. moschata* and suggested that antioxidant activity could depend on the geographical origin of the plants; also, it could result from the deactivation of free radial species by hydrogen atom transfer (HAT). These authors also studied ethanol and methanol *R. sempervirens* and *R. canina* extracts and found that *R. sempervirens* ethanol extracts and *R. canina* methanol extracts were more effective than others in the DPPH assay [37].

Plant extracts intended as human dietary supplements must be non-toxic towards cells. Our studies revealed that the water extracts from leaves and twigs of *R. canina* had no toxic effects on the human kidney cell line HEK 293. Moreover, hemotoxicity was very low, and did not exceed 5% even when the highest concentration was used. Analysis of cytotoxicity to human fibroblasts showed that an ethanolic *R. beggeriana* Schrenk extract was more toxic than the aqueous extract, but both extracts were less toxic to normal than cancer cells [39]. It is known that plant extracts have anticancer potential [48–50]. Importantly, they are harmless to normal cells and do not change their metabolism.

We found that the leaf and twig extracts from *R. canina* did not harm cell membranes, as confirmed by the hemotoxicity assay. Additionally, they were safe for human kidney cells even in high concentrations. Therefore, the tested extracts seem to be appropriate sources of bioactive compounds and antiradical agents for use in the human diet.

*Rosa* plant extracts are considered potent sources of natural antioxidants. It is important to investigate new properties of well-known plants. In our study we have demonstrated antioxidant and antiradical profiles of extracts from *Rosa canina* L. twigs and leaves. Many reports describe rose hips as a valuable source of antioxidants [1–4]. However, numerous tests have revealed that other parts of the plant such as twigs and leaves can also be considered potential sources of beneficial compounds.

## 5. Conclusion

In our study we investigated water extracts from *Rosa canina* leaves and twigs. In general, twig extract was richer in catechins and leaf extract in neochlorogenic acid and ellagic acid. Similar levels of phenolic compounds were found in both. Five B vitamins and three tocopherol isoforms were also found. Also, essential and non-essential amino acids were detected. Circular dichroism revealed that leaf extract interacted more strongly with human serum albumin than twig extract. By checking oxidative stress markers, ROS inhibition and DPPH antiradical scavenging activity, the antioxidant properties of the leaf and twig extracts were revealed. The results suggest that twig extract performed better as an antioxidant. Both extracts were safe for a human kidney cell line and for isolated human erythrocytes. In view of these results it can be concluded that leaf and twig extracts from *Rosa canina* are promising sources of natural compounds, contain valuable nutrient components, and show antiradical effects that should be further investigated.

## Acknowledgments

The authors would like to thank the UK English native speaker (BioMedES Co., UK) for the support in language correction.

